# Auditing Protein-Protein Interaction Signals with Sparse Autoencoder Fingerprints

**DOI:** 10.64898/2026.09.27.754758

**Authors:** Weimin Zhu, Shengfan Wang, Xiaojian Liu, Yiming Xue, Hong-Bin Shen, Badong Chen, Xiaoyong Pan

**Affiliations:** Institute of Image Processing and Pattern Recognition, Shanghai Jiao Tong University, and Key Laboratory of System Control and Information Processing, Ministry of Education of China, Shanghai, 200240, China; State Key Laboratory of Human-Machine Hybrid Augmented Intelligence,Institute of Artificial Intelligence and Robotics,Xi’an Jiaotong University, Xi’an, 710049, China

**Author notes:** Correspondence should be addressed to X. Pan or B. Chen.

## Abstract

Protein language models have become a dominant foundation for sequence-based protein–protein interaction (PPI) prediction, but their generalization remains limited under stringent evaluation, and benchmark accuracy alone cannot reveal whether a PPI predictor learns partner-specific biological signals or exploits contextual shortcuts. Here we introduce AuditPPI, an interpretable framework that transforms sparse-autoencoder (SAE) features from a frozen protein language model into order-invariant pair fingerprints, recasting PPI prediction as an auditable tabular-learning problem. AuditPPI achieves competitive predictive performance across diverse PPI benchmarks while enabling multiscale auditing of the information supporting its predictions. Protein-level audits show that partner-independent participation signals remain substantial in conventional benchmarks, including protein-disjoint benchmark, but are largely insufficient for pair-level discrimination in topology- and degree-controlled benchmark. Pair-level analyses further show that removing protein identity overlap does not eliminate contextual structure: protein-disjoint benchmarking retains strong subcellular co-localization signals and benchmark-dependent feature matching captured by SAE co-activation and absolute-difference. Structural analyses, however, provide little evidence that these predictive signals are specifically grounded in PPI interfaces: feature enrichment at interfaces for protein-disjoint benchmark was no longer detectable after controlling for surface exposure, and contact-specific enrichment was observed for only a small fraction of testable SAE feature pairs. AuditPPI therefore separates predictive success from mechanistic fidelity and provides a practical framework for evaluating whether sequence-based PPI predictors capture partner-specific biological signals instead of contextual shortcuts.

## Main

Protein-protein interactions (PPIs) underpin diverse cellular processes, making their computational prediction a central problem in systems and structural biology^1^. Coevolution-guided structure prediction can resolve residue-level interfaces and complex structures, but often depends on deep paired multiple sequence alignments and intensive computational resources^2,3^. Pretrained protein language models (pLMs)^4,5^ provide a scalable sequence-only alternative, enabling proteins to be represented independently or jointly encoded as pairs for PPI prediction^6–14^. Yet strong benchmark performance does not establish that a model has learned physical binding compatibility. Under conventional pairwise splits, the same proteins can appear in both training and test sets, while close homologues and degree-related network patterns can span the splits. Predictors may therefore exploit protein identity, hub-associated properties or sequence similarity rather than partner-specific interaction signals^15–17^. Negative-sampling strategies introduce additional ambiguity because putative non-interacting pairs are constructed from incomplete interaction annotations and may differ systematically from positive pairs in taxonomy, subcellular localization or broader cellular context^18,19^. These biases are particularly problematic for pLM-based predictors^10^, because pLM representations encode broad biological information^5^, making such contextual differences readily available to downstream classifiers. Consequently, high benchmark accuracy may overstate the extent to which a model has learned transferable, partner-specific biological signals.

Distinguishing partner-specific interaction information from other predictive signals remains difficult because the representations used by pLM-based predictors are high-dimensional and difficult to interpret directly. Recent advances in sparse autoencoders (SAEs)^5,20,21^ provide a way to transform dense pLM representations into sparse feature vocabularies, whose individual features can be examined biologically. Building on this capability, we introduce AuditPPI, an analytical framework that transforms SAE activations from a frozen pLM into order-invariant pair fingerprints while preserving the identities of the underlying SAE features. These fingerprints can be analyzed using standard tabular-learning methods^22–24^. Under matched evaluation protocols, AuditPPI achieves competitive performance across diverse benchmark datasets. The key advantage is that the same SAE vocabulary used for prediction also serves as an audit substrate, allowing pair-level predictive features to be traced back to their corresponding protein- and residue-level SAE features. This design enables partner-independent protein signals, pairwise feature relationships and residue-level structural grounding to be examined within a shared representation framework, and compared across complementary controls for protein reuse, sequence homology, network topology and negative-sampling design. AuditPPI therefore provides a practical framework for separating predictive success from mechanistic fidelity in sequence-based PPI prediction.

## Results

### AuditPPI enables accurate and interpretable PPI prediction

As shown in Fig. 1a, AuditPPI converts sequence-based PPI prediction into an auditable tabular-learning problem. Each protein sequence is encoded using frozen ESMC-6B^5^ and a fixed SAE that defines a 16,384-feature vocabulary with annotations spanning diverse sequence, structural and functional protein properties (Supplementary Fig. S1). Residue-level activations are aggregated via max pooling into protein fingerprints, and two partner fingerprints are combined into the order-invariant pair fingerprint [*A* ∘ *B*, |*A* − *B*|]. Here, *A* ∘ *B* captures feature co-activation between the two proteins, whereas |*A* − *B*| captures differences in feature activation between their fingerprints. The resulting representation supports standard tabular learners while preserving the identities of the underlying SAE features for subsequent auditing.

**Fig 1.**
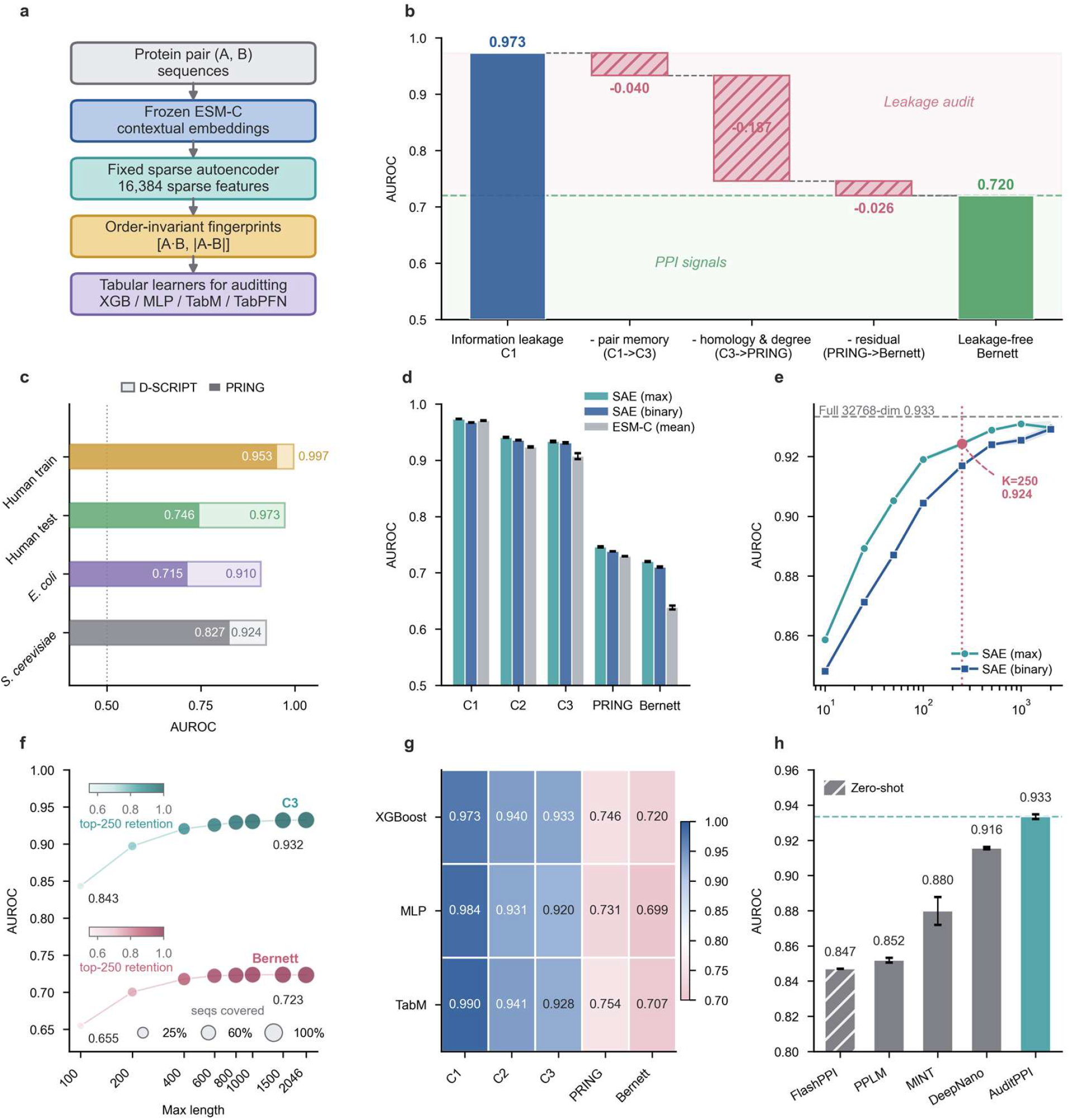
SAE fingerprints enable accurate and auditable sequence-based PPI prediction. **a**, The workflow of AuditPPI. Frozen ESMC-6B representations are passed through a fixed 16,384-feature sparse autoencoder, max-pooled into protein fingerprints and combined into an order-invariant pair fingerprint [*A* ∘ *B*, |*A* − *B*|] for tabular learning. **b**, Test AUROC across C1, C3, the PRING human breadth-first-search split and the Bernett benchmark, representing benchmark settings with increasingly stringent forms of control. **c**, Comparison of lenient D-SCRIPT pair splits and topology-controlled PRING splits in human, *E. coli* and yeast using the same SAE representation and XGBoost learner. **d**, Performance comparison of continuous max-pooled SAE fingerprints, binary SAE-presence fingerprints and mean-pooled ESMC-6B embeddings across benchmark settings. **e**, C3 test AUROC as a function of the number of top-ranked SAE pair features retained. The dashed line indicates performance with the complete pair fingerprint. **f**, Sequence-truncation analysis on C3 and Bernett. Point size indicates the proportion of test proteins fully covered, and color intensity indicates retention of the top 250 SAE features. **g**, Test AUROC of XGBoost, MLP and TabM across five benchmark settings. **h**, Comparison of AuditPPI with DeepNano, MINT, PPLM and FlashPPI on the common C3 benchmark.

We first evaluated AuditPPI across benchmark settings (full dataset statistics are provided in Supplementary Table S1) that impose different types and degrees of control over protein reuse, sequence similarity and graph-related biases (Fig. 1b). Using XGBoost^22^ as the downstream classifier, AuditPPI achieved an area under the receiver operating characteristic curve (AUROC) of 0.973 on C1, where proteins can recur between training and test pairs, and 0.933 on the nominally protein-disjoint C3 split^16^. Performance decreased to 0.746 on the PRING^11^ human breadth-first-search split, which additionally restricts homology and graph-topology reuse, and to 0.720 on the leakage-controlled benchmark constructed by Bernett et al^17^. Because these benchmark families were independently constructed rather than generated by progressively adding controls to the same dataset, differences between them cannot be assigned to individual sources of information reuse. Nevertheless, the above-chance AUROC decreased by 53% between C1 and Bernett, illustrating how performance obtained under conventional pairwise evaluation can fall substantially under more strongly controlled generalization settings.

We next asked whether the contrast between conventional pairwise evaluation and topology-controlled evaluation was consistent across species. Using the same SAE representation and XGBoost learner within each species, we observed the same trend in human, *E. coli* and yeast: replacing the lenient pair split introduced by Sledzieski et al.^6^ with the topology-controlled split PRING^11^ reduced AUROC from 0.973 to 0.746 in human, from 0.910 to 0.715 in *E. coli* and from 0.924 to 0.827 in yeast (Fig. 1c). Although the magnitude of the decrease varied among interactomes, its direction was consistent across all three species. This consistency indicates that benchmark construction strongly influences apparent sequence-based PPI performance and the observed decline is not specific to a single species or interaction network.

Considering that the benchmark design strongly shapes the apparent performance, we further evaluated whether AuditPPI’s sparse representation preserves the predictive information contained in dense ESMC-6B embeddings while making the information directly auditable. As shown in Fig. 1d, max-pooled SAE fingerprints consistently matched or outperformed mean-pooled ESMC-6B representations (layer 60) across benchmark settings, while even a binary SAE-presence fingerprint retained comparable predictive performance.

The sparse representation also allowed us to quantify how concentrated the predictive information was within the SAE vocabulary. Ranking pair-level SAE features by their predictive importance showed that the performance was concentrated in a compact subset: retaining only approximately 250 top-ranked features recovered nearly the full C3 AUROC obtained with the complete 32,768-dimensional pair fingerprint (0.924 versus 0.933; Fig. 1e). Similar feature-concentration analyses across the remaining benchmark settings are provided in Supplementary Fig. S2.

Previous work reported that pLM-based PPI predictors can remain relatively robust to sequence truncation^10^. We therefore asked whether this behavior could be related to the recovery of predictive SAE features. In both C3 and Bernett datasets, AUROC increased and then plateaued as longer sequence segments were retained, closely tracking the recovery of the top 250 SAE features (Fig. 1f). Corresponding sequence-truncation analyses across the remaining benchmark settings are shown in Supplementary Fig. S3. Because the downstream classifier and validation-derived feature ranking were held fixed in this analysis, the loss of performance at shorter sequence lengths was closely associated with incomplete recovery of informative SAE features. Together, these results indicate that SAE fingerprints not only preserve or improve the predictive performance relative to dense ESMC-6B embeddings, but also make the information supporting that performance directly quantifiable.

The benchmark-dependent performance pattern was also reproduced across three downstream learners: XGBoost, a multilayer perceptron (MLP) and TabM (Fig. 1g). Nevertheless, the tabular learners differed systematically in how effectively they used the PPI fingerprints. XGBoost outperformed the MLP on C2, C3, PRING and Bernett, achieving AUROCs of 0.940, 0.933, 0.746 and 0.720, respectively, compared with 0.931, 0.920, 0.731 and 0.699 for the MLP; the MLP was higher only on the least stringent C1 split (0.984 versus 0.973). TabM^24^, which incorporates parameter-efficient ensemble, consistently outperformed the MLP across all five benchmarks, reaching AUROCs of 0.990, 0.941, 0.928, 0.754 and 0.707 on C1, C2, C3, PRING and Bernett, respectively. These results show that AuditPPI is compatible with diverse downstream learners, while tree-based and ensemble tabular methods generally extract the sparse representation more effectively under demanding generalization settings. Additional robustness and control analyses were performed across representations, pLM backbones, sequence lengths, pair-encoding strategies, downstream learners and physicochemical descriptors. Methodological details are provided in Section 4 of Supplementary Methods with the corresponding results shown in Supplementary Tables S2–S6.

Finally, we compared AuditPPI with four adapted sequence-based PPI baselines on common benchmark-specific test pairs. On C3, AuditPPI with XGBoost achieved an AUROC of 0.933. This performance surpassed all adapted sequence-based baselines under the same protocol, including DeepNano^7^ (0.916), MINT^13^ (0.880), PPLM^9^ (0.852) and FlashPPI^12^ (0.847; Fig. 1h). Full benchmark-wise comparisons with the adapted sequence-based baselines are provided in Supplementary Table S7. AuditPPI achieved the highest AUROC and AUPRC in 15 of 16 evaluated settings, with FlashPPI performing the best only on the PRING *E. coli* evaluation.

### Protein-level audit reveals benchmark-dependent participation signals

At the protein scale, we asked two questions: whether a protein’s tendency to participate in positive interactions could be predicted from sequence-derived features, and whether this partner-independent information alone could discriminate PPI labels. Participation was defined according to benchmark structure: positive-network node degree in PRING and labeled-pair participation propensity in C1–C3.

Using the PRING human graph, we defined high-degree proteins as those in the upper decile of the node-degree distribution (degree ≥ 61; Supplementary Fig. S4). An XGBoost classifier using only the SAE fingerprint of each protein predicted high-degree status substantially above chance on the protein-disjoint PRING splits (Fig. 2a and Supplementary Fig. S5). However, assigning each protein pair the lower of its two predicted high-degree probabilities yielded only weak PPI discrimination, with AUROC close to chance. Thus, although high-degree status is predictable from single-protein sequence representations, PRING largely prevents this information alone from determining the pair label.

**Fig 2.**
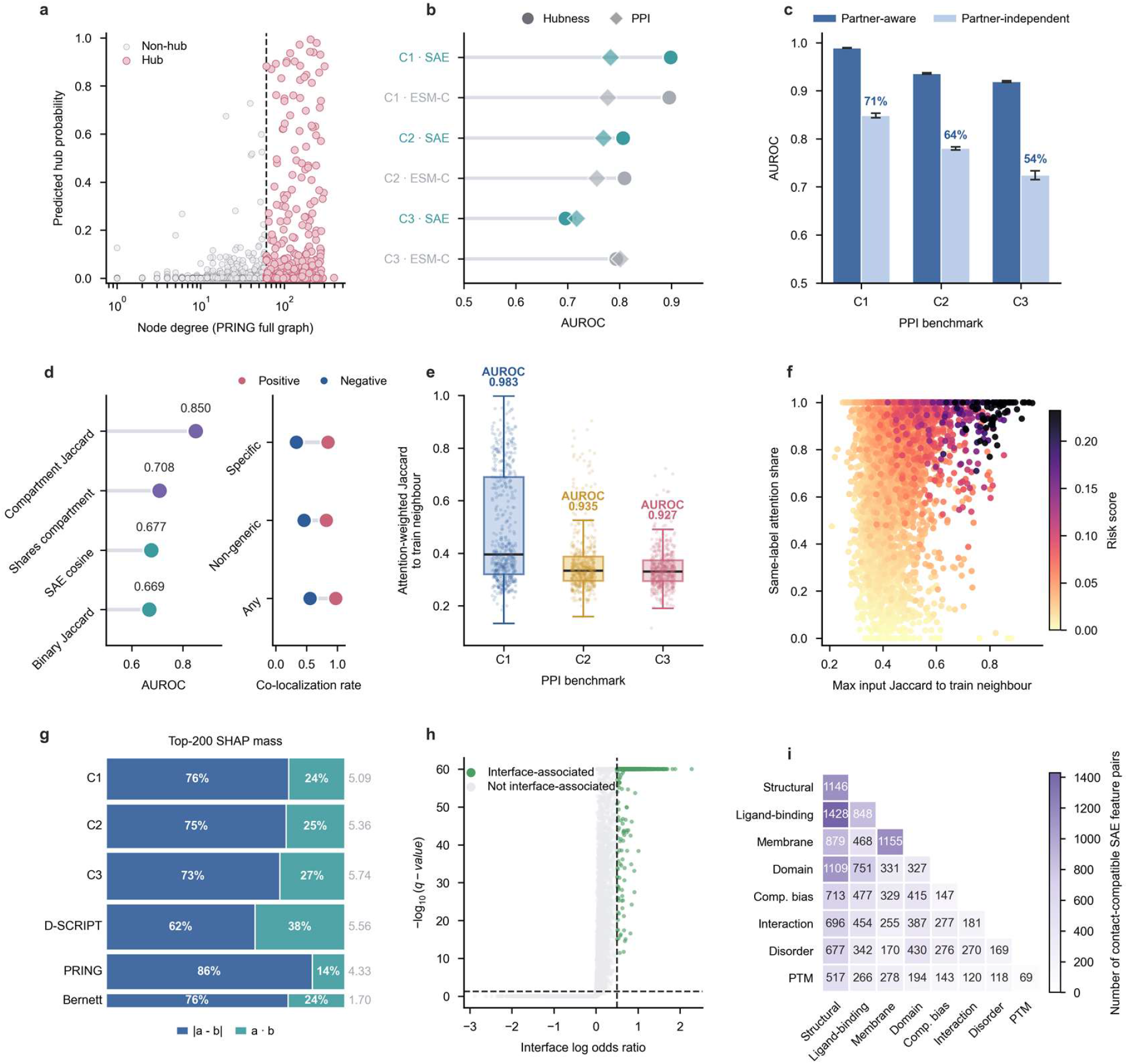
Three-scale auditing separates predictive shortcuts from contact-related signals. **a**, Prediction of high-degree status from single-protein binary SAE fingerprints on the PRING human breadth-first-search split. **b**, Protein-level high-participation prediction, shown as circles, and minimum-participation pair classification, shown as diamonds, on C1, C2 and C3 using SAE or ESMC-6B representations. **c**, Fraction of full pair-model performance recovered by an additive-only model that processes the two proteins independently and contains no pairwise interaction terms. **d**, Model-free discrimination of C3 labels by compartment Jaccard, shared-compartment status, continuous SAE cosine similarity and binary SAE-feature Jaccard. The right panel compares positive- and negative-pair co-localization under increasingly restrictive compartment definitions. **e**, Attention-weighted feature Jaccard similarity between each TabPFN test query and its top training neighbors on C1, C2 and C3. **f**, Retrieval-neighbor audit of C3 test pairs. Each point represents one test query and is colored by a risk score combining maximum feature similarity to an attended training neighbor with the same-label attention share. **g**, XGBoost SHAP attribution across the top 200 SAE pair features, partitioned between the absolute-difference block, |*A* − *B*|, and the co-activation block, *A* ∘ *B*. **h**, Interface association of structurally testable SAE features, shown as log odds ratio for interface enrichment versus Benjamini–Hochberg-adjusted q value; dashed lines indicate the corresponding thresholds. Dashed lines indicate the log-odds-ratio and q-value thresholds. **i**, Annotation-category composition of contact-compatible SAE feature pairs across the eight displayed categories.

TreeSHAP analysis further identified the SAE features associated with the predicted high-degree score. SAE f/6675, annotated as detecting compact ribosomal/RNP-binding cores, was the highest-ranked feature across the breadth-first-search, depth-first-search and random-walk partitions for both binary and max-pooled SAE representations. Additional recurrent high-ranking features were associated with basic extensions and RNA-binding regions of nucleic-acid-associated proteins, including SAE f/5379 and SAE f/16000 in the binary models. Activation of these features was associated with positive contributions to the predicted high-degree score, although the importance of most other features varied across graph partitions. These results identify a recurrent RNA/ribosome-associated feature signature of the fitted high-degree classifiers, but do not imply that these features determine PRING pair-level PPI predictions.

We next applied the analogous audit to the conventional C1–C3 benchmarks. The SAE fingerprint predicted high-participation status with AUROCs of 0.898, 0.807 and 0.695 on C1, C2 and C3, respectively, mean-pooled ESMC-6B embeddings yielded corresponding AUROCs of 0.896, 0.809 and 0.793. We then assigned each protein pair the lower of the two predicted participation probabilities. This minimum-participation score alone separated positive and negative pairs, yielding AUROCs of 0.782, 0.768 and 0.717 on C1, C2 and C3 for the SAE representation, and 0.777, 0.756 and 0.801 for mean-pooled ESMC-6B embeddings. Unlike PRING, the C1–C3 benchmarks therefore retain substantial partner-independent protein-level information, and this signal remains detectable even under the protein-disjoint C3 evaluation.

Consistent with these results, we next asked how much of the full pair model’s performance could be recovered without modelling relationships between the two proteins. A bias-free additive MLP applied the same scoring network to each protein independently and summed the two scores, recovering 71%, 64% and 54% of the full model’s above-chance AUROC on C1, C2 and C3, respectively (Fig. 2c). Because this neural model provides limited direct feature-level interpretation, we additionally fitted a main-effects-only Explainable Boosting Machine (EBM), in which individual SAE features contribute through separate nonlinear shape functions and feature interactions are disabled. This provided an interpretable additive counterpart for examining which SAE features contributed to the partner-independent signal and reproduced the same benchmark-dependent performance pattern (Supplementary Fig. S6). Across additional benchmark settings, species and graph partitions, the extent to which such information could discriminate positive and negative pairs varied strongly with benchmark construction (Supplementary Figs. S5–S6).

EBM feature rankings were highly stable across fitting seeds within C1–C3, with mean pairwise Spearman correlations of 0.989, 0.976 and 0.983, respectively. Several high-ranking features recurred across all three benchmarks, including SAE f/15227, annotated as an NTPase-associated helical element; SAE f/654, associated with extended low-complexity disordered regions; and SAE f/6675, associated with compact ribosomal/RNP-binding cores. Higher activation of SAE f/15227 and SAE f/6675 was associated with more positive fitted EBM contributions, whereas SAE f/654 showed the opposite pattern.

Notably, SAE f/6675 was prominent in both the PRING high-degree classifiers and the C1–C3 additive EBM models, whereas the broader feature rankings differed substantially between the two analyses. This overlap indicates that some sequence-derived protein properties contribute to multiple forms of participation-related information, but the distinct rankings reinforce that positive-network degree in PRING and labeled-pair participation propensity in C1–C3 capture different protein-level signals. Collectively, these analyses suggest that partner-independent information is encoded in interpretable SAE features, while its ability to translate into pair-level PPI discrimination depends strongly on benchmark construction.

### Pair-level audit reveals contextual concordance and feature-level matching signals

Having identified partner-independent participation signals at the protein level, we next asked whether simple relationships between the two protein representations could discriminate PPI labels without fitting a pair classifier. We first focused on C3, the protein-disjoint benchmark in which such pair-level signals cannot arise from direct reuse of individual proteins across train and test sets. The cosine similarity between continuous SAE fingerprints and the Jaccard similarity between binary SAE features yielded AUROCs of 0.677 and 0.669, respectively (Fig. 2d). Subcellular localization provided a substantially stronger contextual signal: the Jaccard overlap between the two proteins’ annotated compartment sets reached an AUROC of 0.850, and 97.4% of positive pairs shared at least one compartment compared with 55.8% of negative pairs. This difference remained pronounced when the analysis was restricted to non-ubiquitous compartments, with co-localization rates of 84.6% and 33.2% for positive and negative pairs, respectively.

To assess whether SAE similarity remained informative after accounting for this coarse localization signal, C3 test pairs were stratified according to whether the two proteins shared at least one annotated compartment, and SAE cosine similarity was evaluated separately within each subset. SAE cosine remained predictive among pairs already sharing at least one compartment (AUROC 0.655; n = 3,683) and was also informative among non-co-localized pairs (AUROC 0.760). Under progressively more restrictive compartment definitions, the AUROC within co-localized subsets remained above chance (0.655, 0.716 and 0.724 for all compartments, exclusion of nucleus and cytoplasm, and the predefined non-ubiquitous subset, respectively). Thus, subcellular co-localization is a strong contextual signal in C3 but does not fully account for the information captured by SAE representation similarity.

Beyond these model-free measures of pair similarity, we next examined whether held-out protein pairs had closely related labeled examples in the training set. Using TabPFN as a post-hoc diagnostic, we identified the labeled training pairs receiving the highest decoder attention from each test pair and measured their overlap in the reduced SAE pair-feature representation. Across C1–C3, highly attended training pairs showed feature overlap with their corresponding test pairs, with the strongest overlap observed in C1 (Fig. 2e). On C3, a median of 85% of the attention mass within the top attended neighborhood was assigned to training pairs carrying the same label as the test pair. Using the prespecified thresholds described in Methods, 1.2% of C3 test pairs met the high-risk retrieval criterion, combining high pair-feature overlap with strongly concentrated attention on same-label training examples (Fig. 2f). These results reveal label-concordant neighborhood structure in the C3 SAE representation space, but are interpreted as a post-hoc diagnostic rather than a causal explanation of the primary XGBoost predictions. Similar labeled-neighbor structure was observed across the other benchmark settings, although its prevalence and label concordance varied substantially with the evaluation regime (Supplementary Fig. S7).

Feature attribution next showed that predictive importance was concentrated in the absolute-difference block of the pair representation. Among the top 200 SAE pair features ranked by mean absolute TreeSHAP attribution, the absolute-difference block accounted for 73%, 76% and 86% of the summed attribution on C3, Bernett and PRING, respectively, whereas co-activation contributed more strongly in the cross-species setting (Fig. 2g). Importantly, dominance of the absolute-difference block does not imply that greater feature discordance favored positive predictions. Feature-level SHAP profiles showed that, for many of the highest-ranking absolute-difference features, increasing absolute-difference instead shifted the model towards the negative class. Thus, these models frequently used absolute-difference variables as measures of feature-wise matching rather than as evidence that dissimilar proteins were more likely to interact.

We next examined the annotation-based biological interpretation of the SAE pair features used by the XGBoost models. Across benchmark settings, PTM-labeled and disorder-associated SAE features were consistently over-represented among the top-ranked pair features relative to their representation in the full SAE codebook, although both category composition and the balance between co-activation and absolute-difference features remained benchmark dependent (Supplementary Fig. S8). The PTM annotation category in this resource includes many features associated with signal peptides, organelle-targeting peptides and protein processing or maturation, and therefore should not be interpreted specifically as enrichment of phosphorylation-related features.

Feature-level TreeSHAP profiles further revealed distinct biological themes in C3 and Bernett (Supplementary Fig. S9). In C3, the highest-ranking pair feature was the absolute difference in SAE f/7262, annotated as detecting the N-terminal and hydrophobic regions of secretory/ER-targeting signal peptides. Other highly ranked features included co-activation of SAE f/6675, associated with compact ribosomal/RNP-binding cores, and the absolute difference in the signal-peptide-associated SAE f/15889. Larger values of the SAE 7262 and SAE 15889 absolute-difference features were associated with more negative model contributions, whereas stronger co-activation of SAE f/6675 was associated with more positive contributions. Several additional high-ranking co-activation features, including SAE f/15318, SAE f/7150 and SAE f/5530, were associated with organelle-or mitochondrial-targeting peptides and likewise contributed positively with increasing co-activation. Together, these patterns indicate that the C3 classifier exploits concordance in secretory-pathway-associated properties together with shared activation of RNA/RNP- and organelle-targeting-associated features.

Notably, SAE f/6675 also ranked prominently in the protein-level participation analyses. Its interpretation differs between the two scales: the protein-level models associate the marginal activation of SAE f/6675 in an individual protein with participation propensity, whereas the C3 pair model uses the joint activation of this feature in both proteins. Its recurrence therefore links protein-level and pair-level signals without implying that the same predictive mechanism operates in the two analyses.

Bernett showed a different feature profile, with mitochondrial-targeting and membrane-associated features particularly prominent. The strongest pair feature was co-activation of SAE f/5530, associated with mitochondrial targeting presequences and related amphipathic basic N termini. Membrane-associated SAE dimensions also entered through both pair transformations. For example, SAE f/5659, associated with hydrophobic transmembrane helices, appeared as both a co-activation and an absolute-difference feature: greater co-activation was associated with positive model contributions, whereas increasing absolute-difference was associated with negative contributions. More generally, individual SAE dimensions could enter the models through different pair relationships across benchmarks; for example, SAE f/7262 appeared as an absolute-difference feature in C3 but as a co-activation feature in Bernett. These observations show that biological annotation alone does not determine a feature’s predictive role: the relevant signal depends on how activation of that latent relates across the two proteins and on the benchmark in which it is evaluated. These analyses provide an annotation-based interpretation of the biological information exploited by the fitted XGBoost pair models, but they do not establish that the corresponding sequence features are physically localized to protein–protein interaction interfaces. We therefore next tested their structural grounding at residue resolution.

### Residue-level audit distinguishes surface-associated predictive signals from contacts

At the residue scale, we asked whether SAE features ranked as important for C3 prediction were specifically associated with protein–protein interfaces. Using non-homologous PDB complexes, 11.1% of structurally testable SAE features were significantly enriched at interface residues, and the top 50 C3-ranked features were enriched 1.98-fold among this interface-associated set (P = 0.020; Fig. 2h). However, when the control was restricted to solvent-exposed non-interface residues, this enrichment decreased to 1.12-fold and was no longer significant (P = 0.435). Thus, the apparent interface enrichment of C3-predictive SAE features was largely explained by general surface exposure rather than interface specificity.

We therefore applied a stricter residue-pair test of contact compatibility, comparing SAE feature-pair co-activation across cross-chain contacts (C_β_–C_β_ distance ≤ 8 Å) with spatially distant controls (C_β_–C_β_ distance > 12 Å). Contact-enriched feature pairs spanned several SAE annotation categories, including structural motifs, ligand-binding sites and membrane-associated regions (Fig. 2i). These results indicate that the SAE representation contains contact-related structural information, even though the C3-predictive single-feature signals show little evidence of broad interface specificity after controlling for surface exposure.

## Discussion

AuditPPI shows that strong sequence-based PPI performance can coexist with limited evidence that a predictor has learned broadly transferable, partner-specific physical compatibility. Removing protein overlap between training and test sets, as in C3, does not eliminate other predictive information: partner-independent participation signals remain detectable, and positive and negative pairs differ strongly in subcellular co-localization. Pair-level feature attribution further shows that the model exploits structured relationships between the two protein representations. In C3, many highly ranked absolute-difference features contributed more negatively as the two proteins became more dissimilar, while several co-activation features contributed positively when both proteins activated the same sequence-derived property. These patterns indicate that absolute-difference variables often function as measures of feature matching rather than evidence that greater feature discordance favors interaction. The biological content and pairwise form of these signals were also benchmark dependent, with secretory-, organelle-targeting- and RNA/RNP-associated features prominent in C3 and mitochondrial- and membrane-associated features more prominent in Bernett. These predictive relationships should not, however, be equated with physical interaction mechanisms. The C3 SAE representation space contains feature-similar, label-concordant training examples, but this neighborhood structure is a post-hoc property of the representation and does not establish what causally drives the primary XGBoost predictions. Likewise, residue-level analyses provided only limited evidence that predictive SAE signals are directly grounded at protein interfaces. Features ranked as important for C3 prediction appeared enriched at interface residues, but this enrichment was no longer detectable after comparison with solvent-exposed non-interface residues. In a separate residue-pair analysis, only a minority of testable SAE feature pairs were enriched among contacting relative to spatially distant residue pairs. The SAE vocabulary therefore contains contact-related structural information, but the current results do not establish that this information is sufficiently widespread, or sufficiently aligned with the features driving benchmark prediction, to explain overall predictive performance.

AuditPPI is diagnostic rather than corrective. It identifies which protein-level, pair-level and structural signals are available to a predictor, but does not itself remove these signals from the data or force a model to rely on partner-specific physical compatibility. The audits nevertheless suggest practical directions for benchmark design. Protein-disjoint and sequence-redundancy-controlled splits can limit protein identity and homology reuse, while degree-or participation-aware sampling can reduce the extent to which single-protein propensity alone discriminates pair labels. Negative sampling could additionally be matched for broad biological context, including subcellular localization or taxonomy, so that such properties alone are insufficient to separate positive from negative pairs. Pair-level matching signals may require further controls that balance biologically relevant properties across classes without erasing the interaction task itself. This distinction is important because cellular context is genuinely informative about whether two proteins can encounter and interact; removing it too aggressively may create an artificial prediction problem or increase false-negative contamination. The goal of benchmark control should therefore be to prevent contextual compatibility from becoming sufficient for prediction, rather than to eliminate biologically meaningful context altogether.

Several limitations remain. First, most AuditPPI analyses measure the availability, predictive sufficiency or fitted contribution of candidate signals rather than causally decomposing individual predictions. TreeSHAP describes how a fitted model uses pair features, but does not establish that the corresponding biological property causally determines interaction. Second, biological interpretation depends on the chosen SAE dictionary and the associated feature annotations, and annotation categories can be broad; for example, features grouped under post-translational modification include signal-peptide, organelle-targeting and protein-processing signals rather than representing a single molecular mechanism. Finally, structural analyses are necessarily biased towards experimentally resolved complexes and may underrepresent transient, disordered and condition-dependent interactions. Future work could combine audit-guided benchmark construction with causal feature interventions, context-matched evaluation and broader structural or experimental data to test whether reducing benchmark-associated predictive signals leads models to rely more directly on partner-specific interaction information.

## Methods

### Overview of AuditPPI

AuditPPI is a predictive and diagnostic framework that converts sequence-based PPI prediction into an auditable tabular-learning problem. Each protein sequence was encoded using the frozen ESMC-6B protein language model, and residue-level representations extracted from layer 60 were passed to a fixed sparse autoencoder trained specifically on layer-60 ESMC-6B representations. The resulting sparse features were aggregated into protein-level fingerprints and combined into an order-invariant pair-level representation for downstream classification.

For a protein *p* with an original sequence length *L*_p_, the effective input length was defined as 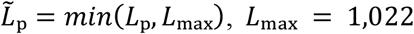. Proteins longer than 1,022 residues were truncated before ESMC-6B encoding. After excluding special tokens, the frozen protein language model produced a contextual residue-embedding matrix:

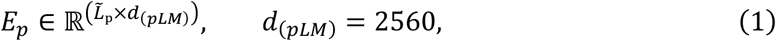

where *d*_(pLM)_ denotes the dimensionality of the ESMC-6B residue embeddings. The frozen SAE encoder subsequently mapped the dense residue embeddings to sparse feature activations:

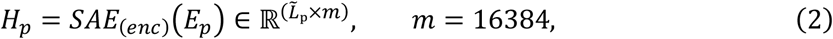

where *H*_p,r,k_ denotes the activation of SAE feature *k* at residue *r* of protein *p*. The SAE used a TopK activation constraint with *K* = 64, retaining up to 64 non-zero SAE feature activations at each residue. A continuous protein-level fingerprint was generated by max pooling each SAE feature over all retained residues:

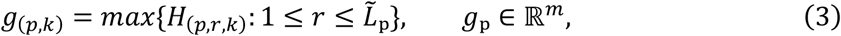

and a binary fingerprint

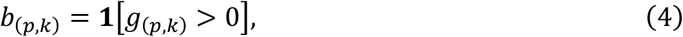

Thus, the continuous fingerprint preserved the magnitude of feature activation, whereas the binary fingerprint recorded only whether a feature was activated at least once in the protein.

For proteins a and b, their protein fingerprints were combined into an order-invariant pair fingerprint:

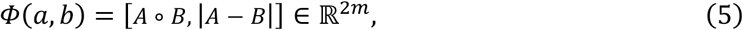

where *A*/*B* denotes either the continuous fingerprint *g*_p_ or the binary fingerprint *b*_p_, and ∘ denotes element-wise multiplication. The first block captures feature co-activation between the two proteins, whereas the second captures the absolute difference in feature activation. The resulting pair fingerprints were fed to downstream tabular classifiers, while the shared 16,384-feature SAE vocabulary was retained for subsequent protein-, pair- and residue-level auditing.

Further discussion of the conceptual framework and interpretive scope is provided in Supplementary Note 1, with additional details on the SAE representation provided in Supplementary Note 2.

### PPI datasets and benchmark settings

AuditPPI was evaluated in several benchmarks drawn from four source families: the C1-C3 dataset released by Szymborski and Emad^10^, the D-SCRIPT cross-species benchmark^6^, the PRING graph benchmark^11^ and the Bernett gold-standard dataset^17^. These settings differ in their interaction sources, negative-sampling procedures and generalization objectives. Detailed dataset provenance, partitioning criteria and preprocessing procedures are provided in Section 2 of Supplementary Methods. Full dataset statistics are provided in Supplementary Table S1.

### Tabular learners and performance evaluation

AuditPPI was evaluated using XGBoost, MLP, TabM and TabPFN. XGBoost, the MLP and TabM were used for the principal PPI prediction experiments, providing tree-based, conventional neural and parameter-efficient ensemble learners for the same order-invariant protein-pair representation. Unless otherwise stated, the primary AuditPPI configuration used XGBoost with continuous max-pooled SAE fingerprints and the pair representation [*A* ∘ *B*, |*A* − *B*|]. TabPFN was used for the labeled-neighbor audit described below rather than for the principal benchmark comparisons.

Models were trained independently for each benchmark using its predefined training, validation and test partitions. Early stopping and model selection were performed using validation data only. All experiments were repeated using three random seeds (42, 43 and 44), and the checkpoint or boosting iteration with the highest validation AUROC was used for test evaluation. Performance was assessed primarily using AUROC, with AUPRC additionally reported where class imbalance made precision–recall performance informative. Results are reported as the mean and standard deviation across seeds unless otherwise stated. Detailed implementation settings, software versions and the computational environment are provided in the Supplementary Methods (Section 1,3).

### Sequence-based PPI baselines

We compared AuditPPI with four published sequence-based PPI predictors: DeepNano-seq^7^, MINT^13^, PPLM^9^ and FlashPPI^12^. Frozen pretrained encoders, released checkpoints and author-provided inference implementations were used. Trainable prediction heads were fitted using the training partition of each benchmark family and selected using its validation partition. When no validation partition was supplied, 10% of the training pairs were reserved by stratified sampling. For computational consistency, trainable baseline heads used stratified training subsets of at most 100,000 pairs. Evaluation used the same test pairs, labels, AUROC and AUPRC calculations, and random seeds as AuditPPI. Detailed model-specific adaptation, feature-extraction and scoring procedures, together with the scope of each adapted baseline, are provided in Supplementary Methods, Section 5.

### Protein-level audit of partner-independent interaction propensity

We used a protein-level audit to test whether properties of each protein alone could predict its tendency to participate in positive interactions, and whether this information could in turn discriminate positive from negative protein pairs. Because the benchmark structures differ, participation was defined differently for C1–C3 and PRING. In C1–C3, participation was defined from the released labeled pair tables, which contain both positive interactions and sampled negative pairs. In PRING, participation was defined from positive-network node degree.

For a C1–C3 partition containing labeled protein pairs (*A*_i_, *B*_i_, *y*_i_), where *y*_i_ ∈ {0,1}, we defined the number of labeled pairs containing protein *p* as

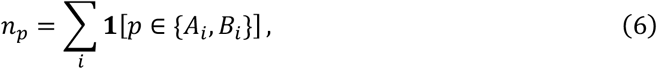

and the number of positive labeled pairs containing *p* as

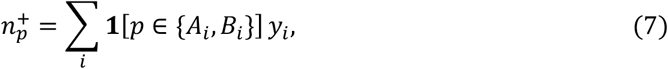

The participation rate of protein *p* was then

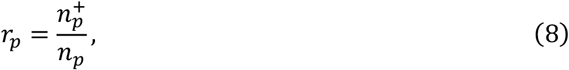

Thus, *r*_p_ measures the fraction of observed labeled pairs involving *p* that are positive. Because *n*_p_ counts both positive and sampled negative pairs, it should not be interpreted as interaction-network degree. These quantities were calculated separately within the training, validation and test partitions.

Let τ_90_ denote the 90th percentile of the training-set participation-rate distribution. A protein was assigned the high-participation audit label

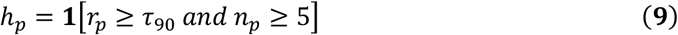

The threshold τ_90_ estimated from the training partition was applied unchanged to validation and test proteins. For the PRING human analysis, the audit label was instead based on positive-network degree *d*_p_. Proteins in the upper decile of the degree distribution were classified as high-degree, corresponding to *d*_p_ ≥ 61.

XGBoost classifiers were trained to predict these benchmark-specific protein-level audit labels from a single-protein representation. Binary SAE-presence fingerprints, continuous max-pooled SAE fingerprints and mean-pooled ESM-C embeddings were evaluated separately. Protein-level performance was measured using AUROC and AUPRC. In C1 and C2, proteins may occur in more than one pair partition under the original benchmark construction; C3 and the PRING breadth-first-search analysis provided protein-disjoint evaluations.

To determine whether the predicted protein-level signal alone could discriminate PPI labels, we converted the two single-protein predictions into a pair score. Let 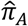 denote the predicted probability that protein *p* belongs to the corresponding high-participation or high-degree audit class. For proteins A and B, we defined the minimum-participation score as

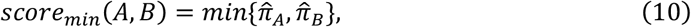

This score is high only when both proteins are independently predicted to have high participation. Because no relationship between the two protein representations is modelled, its AUROC against the PPI labels quantifies how well partner-independent protein-level information alone separates positive and negative pairs.

To estimate how much predictive performance could be obtained from the two proteins independently, we additionally constructed additive protein-only models to estimate how much of the full pair model’s performance could be recovered without using any jointly constructed pair features. For a protein fingerprint *x*_p_, the same scoring function *f* was applied independently to both proteins, and the resulting contributions were summed:

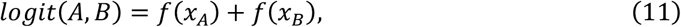

where *logit*(*A, B*) denotes the additive pair logit. For the MLP implementation, *f* was a shared bias-free neural network. For the EBM, *f* consisted only of additive main effects over individual SAE features, with feature-interaction terms disabled. Neither model received feature co-activation, absolute differences or any other jointly constructed pair features. Finally, we defined the above-chance AUROC recovery ratio of an additive model as

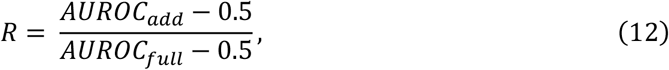

where*AUROC*_add_ is the AUROC of the additive protein-only model and *AUROC*_full_ is the AUROC of the corresponding full AuditPPI pair model. Values were reported as 100*R*%, such that 0% indicates chance-level additive performance and 100% indicates recovery of the full model’s above-chance AUROC.

### Pair-level audit of contextual and neighbor support

We first tested whether positive and negative protein pairs could be distinguished by simple similarity measures without fitting a pair classifier. For each pair, cosine similarity was calculated between the continuous max-pooled SAE fingerprints of the two proteins, and Jaccard similarity was calculated between their sets of active binary SAE features. Both scores were evaluated directly against the C3 interaction labels using AUROC. Detailed definitions are provided in Supplementary Methods, Section 6.1.

We next examined subcellular co-localization as a source of biological-context similarity. UniProt annotations were collected for proteins in the C3 benchmark and mapped to a predefined set of coarse cellular compartments. Gene Ontology Cellular Component annotations were used as the primary source, with UniProt free-text subcellular-localization annotations used when structured annotations were unavailable. For each protein pair, we recorded whether the two proteins shared at least one compartment and calculated the Jaccard similarity between their compartment sets. These scores were evaluated directly against the C3 test labels using co-localization rates and AUROC, without fitting a classifier.

Robustness was assessed using three compartment definitions: all mapped compartments, exclusion of the broadly distributed nucleus and cytoplasm categories, and a predefined subset of non-ubiquitous compartments. To determine whether SAE similarity remained informative after accounting for coarse localization, C3 test pairs were stratified according to whether the two proteins shared at least one compartment under each definition. Cosine similarity between their continuous SAE fingerprints was then evaluated separately within the co-localized and non-co-localized subsets. Detailed annotation mapping, filtering rules and statistical procedures are provided in Supplementary Methods, Section 6.2 and Supplementary Tables S8–S9.

We additionally used TabPFN as a post-hoc diagnostic to determine whether held-out protein pairs had feature-similar labeled examples in the training set. For each benchmark family, XGBoost was first fitted to the complete 32,768-dimensional SAE pair representation, and pair columns were ranked by mean absolute TreeSHAP attribution on the corresponding validation partition. Each SAE identifier inherited the better rank of its co-activation and absolute-difference columns. The 200 highest-ranked identifiers were retained together with both associated pair columns, yielding a 400-dimensional representation for the TabPFN analysis.

For each held-out query pair *i*, decoder attention weights to the labeled training pairs were extracted from the fitted TabPFN model. The top *K* = 20 attended training pairs, *N*_K_(*i*), were retained. Similarity between two reduced pair fingerprints was calculated as Jaccard overlap between their nonzero co-activation and absolute-difference columns.

The attention-weighted neighbor overlap was calculated as:

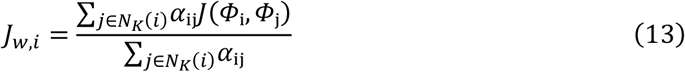

where α_ij_ denotes the attention weight from query *i* to training pair *j*. Label-concordant support was calculated as:

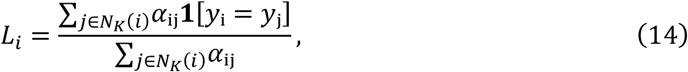

The retrieval-risk score was defined as:

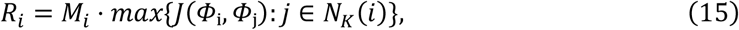

where *M*_i_ = ∑_{∈ K(i),yj= yi}_*α*_i_ for the same-label attention mass. High-risk pairs were identified using the prespecified thresholds on retrieval-risk score, maximum feature overlap and same-label attention share reported in the Supplementary Methods (Section 6.3).

The relative use of pair concordance and discordance was quantified using the validation-derived TreeSHAP values described in “Tabular learners and performance evaluation”. Attribution was summed separately over the co-activation and absolute-difference blocks among the selected feature columns and normalized by their combined attribution. The biological composition of highly attributed pair variables was analyzed by mapping their underlying SAE identifiers to the released ESMC-6B annotation taxonomy and comparing category frequencies with the complete SAE codebook. Cross-benchmark category enrichment, block-specific composition and feature-level TreeSHAP interpretation are described in Supplementary Methods, Section 7.

### Residue-level audit of structural grounding

Residue-level analyses used non-homologous positive complexes from the PDB structural dataset. SAE activations were generated directly from structure-derived chain sequences, preserving correspondence between encoded positions and resolved structural residues by construction. Details of structure-derived sequence construction, residue indexing, coordinate assignment and sequence-length reconciliation are provided in Supplementary Methods, Section 8.1.

For two interacting chains A and B, residue r in chain A was classified as an interface residue when at least one residue in chain B was within a C_β_–C_β_ distance of 8 Å:

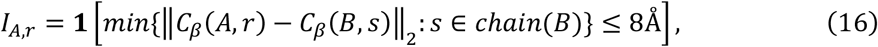

The same definition was applied symmetrically to chain B. SAE feature *k* was considered active at a residue when its cached activation at the corresponding sequence position was greater than zero.

We first tested whether individual SAE features were preferentially associated with protein– protein interfaces. Two non-interface backgrounds were considered. The broad background contained all non-interface residues from the same analyzed chains. The surface-matched background contained only solvent-exposed non-interface residues, with solvent exposure defined as relative solvent-accessible surface area *rSASA* ≥ 0.20. Solvent-accessibility calculations and the exact eligibility criteria used to define structurally testable SAE features are provided in Supplementary Methods, Section 8.2.

For each structurally testable SAE feature k, active and inactive residues were counted separately in the interface and control sets. Let 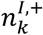 and 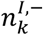 denote the numbers of active and inactive interface residues, respectively, and 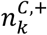 and 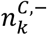 the corresponding numbers in the control background. Interface enrichment was summarized by the odds ratio

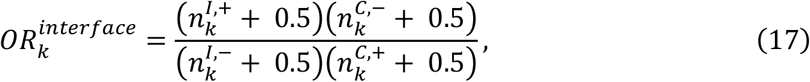

where the 0.5 pseudocount corresponds to the Haldane–Anscombe correction. Statistical significance was assessed on the corresponding contingency tables using one-sided Fisher’s exact tests followed by Benjamini-Hochberg correction across tested SAE features. A feature was classified as interface-associated when 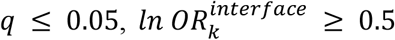, it was active in at least 25 interface-residue instances and its interface occurrences were supported by at least five distinct complexes.

We subsequently whether examined SAE features important for C3 prediction were over-represented among the interface-associated features. For this structural audit, validation-derived TreeSHAP attribution from the co-activation and absolute-difference columns corresponding to the same SAE identifier was combined to obtain an identifier-level C3 ranking. The top 50 C3-ranked SAE identifiers were then tested for enrichment among interface-associated features using a hypergeometric test. The background universe was restricted to SAE identifiers that were structurally testable under the corresponding control definition rather than to the complete SAE vocabulary. This analysis was performed independently using the broad and surface-matched backgrounds, allowing apparent interface enrichment of C3-predictive features to be reassessed after controlling for general surface exposure. Additional implementation details are provided in Supplementary Methods, Section 8.2.

Independently of the C3 predictive-feature ranking, we next tested whether the SAE vocabulary contained feature-pair associations enriched at physical cross-chain contacts. Cross-chain residue pairs with C_β_–C_β_ distance ≤ 8 Å formed the contact set. Surface-exposed, non-interface residue pairs from the same complex with C_β_–C_β_ distance > 12 Å formed the control set and were sampled within each complex to approximately match the number of contact pairs. For each retained residue, the four SAE features with the largest positive activation values were selected to represent its dominant local sparse features. For each cross-chain residue pair, the Cartesian product of the two four-feature sets generated up to 16 feature combinations. Feature combinations were treated as unordered SAE-identifier pairs, and duplicate occurrences of the same unordered feature pair within a residue pair were counted only once.

For each structurally testable SAE feature pair *u*, let 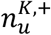 and 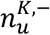 denote the numbers of contact residue pairs in which *u* was present or absent, respectively, and 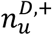 and 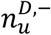 the corresponding counts among distant-control residue pairs. Contact enrichment was summarized as

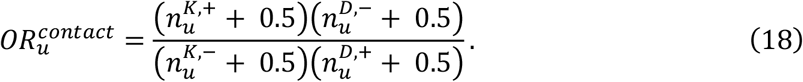

Feature-pair enrichment at contacts was assessed using one-sided Fisher’s exact tests followed by Benjamini-Hochberg correction across tested feature pairs. A feature pair was classified as contact-compatible when 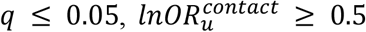, it occurred in at least 100 contact residue-pair instances and its contact occurrences were supported by at least ten distinct complexes. Biological annotations were assigned only after statistical classification of contact-compatible feature pairs and were used for descriptive interpretation. Detailed annotation and category-level aggregation procedures are provided in Supplementary Methods, Section 8.3.

## Data availability

All datasets used in this paper are publicly available. The C1-C3 datasets are available at https://zenodo.org/records/17353963. The D-SCRIPT’s cross-species dataset created by Sledzieski et al. is available at https://d-script.readthedocs.io/en/stable/data.html. The PRING dataset is available at https://github.com/SophieSarceau/PRING. The human dataset created by Bernett et al. is available at https://doi.org/10.6084/m9.figshare.21591618.v3. The non-homologous PDB protein–protein complexes used for the residue-level audit were obtained from Zhang et al.^3^, which is available at https://conglab.swmed.edu/humanPPI/humanPPI_download.html. The archived version of these datasets is available via Zenodo at https://zenodo.org/uploads/21512494. Source data are provided with this paper.

## Code availability

The source code used in this study is available at https://github.com/jaminzzz/AuditPPI under the MIT license. The specific version of the code associated with this publication is archived in Zenodo and is accessible via https://zenodo.org/uploads/21512494.

## Acknowledgements

This work is supported by the National Natural Science Foundation of China (No. 62473257, 62573293, 32541016), and the Science and Technology Commission of Shanghai Municipality (24ZR1435300, 24510714300).

## Author contributions

X.P. and B.C. conceived and supervised the project. W.Z. collected and preprocessed the data and developed and trained the model under the methodological guidance of X.P. and B.C., W.Z., S.W., X.L., Y.X., and X.P. drafted the manuscript, with contributions from H.B.S. All authors reviewed, revised, and approved the final version of the manuscript.

